# Streptococcal superantigen-induced expansion of human tonsil T cells leads to altered T follicular helper cell phenotype, B cell death, and reduced immunoglobulin release

**DOI:** 10.1101/395608

**Authors:** Frances J. Davies, Carl Olme, Nicola N. Lynskey, Claire E. Turner, Shiranee Sriskandan

## Abstract

**Background:** Streptococcal pyrogenic exotoxin (SPE)A expression is epidemiologically linked to streptococcal tonsillo-pharyngitis and outbreaks of scarlet fever, although the mechanisms by which superantigens confer advantage *Streptococcus pyogenes* are unclear. *S. pyogenes* is an exclusively human pathogen. As the leukocyte profile of tonsil differs from peripheral blood, the impact of SPEA production on human tonsil cell function was investigated.

**Methods:** Human tonsil cells from routine tonsillectomy, were co-incubated with purified streptococcal superantigens or isogenic streptococcal culture supernatants, differing only in superantigen content. Tonsil cell proliferation was quantified by tritium-incorporation, and cell surface characteristics assessed by flow-cytometry. Soluble mediators were measured using ELISA and quantitative (q)RT-PCR was performed for immunoglobulin gene expression.

**Results:** Tonsil T cells proliferated in response to SPEA and demonstrated typical release of pro-inflammatory cytokines. When cultured in the absence of superantigen, tonsil preparations released large quantities of immunoglobulin over 7d. In contrast, marked B cell apoptosis and abrogation of total IgA, IgM, and IgG production occurred in the presence of SPEA and other superantigens. In SPEA-stimulated cultures, T follicular helper (TfH) cells showed a reduction in CXCR5 expression, but up-regulation of CD134 (OX40), CD278 (ICOS) and CD150 (SLAM) expression, indicative of a phenotypic change in the TfH population and associated with impaired chemotactic response to CXCL13.

**Conclusions:** SPEA and other superantigens cause dysregulated tonsil immune function, driving T cells from TFH to a proliferating phenotype, with resultant loss of B cells and immunoglobulin production, providing superantigen-producing bacteria with a likely survival advantage.

## Introduction

The virulent human pathogen *Streptococcus pyogenes* can produce up to 11 different secreted superantigens that contribute to the features of cytokine-induced toxic shock during lethal, invasive infections such as necrotising fasciitis (Proft & Fraser, 2016). Invasive infections are, however, rare compared with symptomatic non-invasive disease that occurs in the nasopharynx, manifest as pharyngitis, tonsillitis, and the childhood exanthem scarlet fever. Indeed, in human populations, the nasopharynx represents the main reservoir of *S. pyogenes*.

Upon secretion in the vicinity of host leukocytes, streptococcal superantigens bind host MHCII outside the antigen groove and ligate a variably discrete repertoire of T cell receptor variable β chain (TCRVβ), thereby leading to mass activation and proliferation of all target populations of T cells that bear relevant TCRVβ (Faulkner et al., 2005). As such, the evolutionary benefit of superantigen production is most likely conferred to *S. pyogenes* through activation of T cells within the nasopharynx and, in particular, the human tonsil, in ways that provide a survival or transmission advantage. The tonsil is a solid secondary lymphoid organ that possesses only efferent lymphatic drainage; the leukocyte populations that constitute the tonsil are distinct from those present in peripheral blood and also distinct from mucosal lymphoid tissue, and comprise follicular dendritic cells, T follicular helper (Tfh) cells, and B cells arranged in germinal centres, bounded by the specialised tonsil mucosal epithelium in the posterior nasopharynx (Jovic et al., 2015).

Streptococcal expression of superantigen genes is increased upon exposure to tonsil epithelium (Broudy, et al., 2001) and in non-human primate models of tonsillo-pharyngitis (Virtaneva et al., 2005). Furthermore, the phage-encoded superantigen streptococcal pyrogenic exotoxin A (SpeA) is required for successful nasopharyngeal infection with *S. pyogenes* in HLA-DQ8 superantigen-sensitive transgenic mice, albeit that mice lack tonsils (Kasper et al., 2014). It is therefore remarkable that, in contrast to work that has elucidated effects of superantigens on rodent spleen cells and leukocytes of human peripheral blood (Proft & Fraser, 2016), the impact of superantigens on human tonsil has not been addressed. Using the phage-encoded toxin SpeA, that has been implicated in the emergence of severe *emm*1 *S. pyogenes* infections in recent decades, as a paradigm of a streptococcal superantigen (Stevens et al., 1989) we set out to conduct a systematic examination of responses to superantigen using both human tonsil histocultures and cultured cell suspensions.

## Methods

### Reagents

Recombinant toxins were purchased from Toxin Technology, Sarasota Florida. Recombinant SMEZ and SPEJ were a kind gift of Thomas Proft (Auckland, NZ), produced as previously described (Proft, et al., 1999). Concanavalin A (Sigma), anti-CD3 and anti-CD28 (Milltenyi Biotech) were used at 1 μg/ml per supplier recommendations. Antibodies for flow cytometry (Supplementary Table 1) were purchased from BD Biosciences, conjugated to either FITC, PE, PECy5 or PerCPCy7, with the following exceptions: TCRVβ antibodies, isotype controls and TCRVβ Repertoire kit (IOTest Beta Mark) Beckman Coulter (Marseille, France), CXCR5 PE and isotype control R&D systems, CXCR5 PE-Cy5 and isotype control eBioscience (California, USA).

### Bacterial strains

Isogenic *emm*1/M1 *S. pyogenes* strains that differ in SPEA production (Sriskandan et al., 2001) or isogenic *emm*89/M89 strains that differ in SMEZ production (Unnikrishnan et al., 2002) were cultured overnight in RPMI supplemented with 10% FBS and cell free supernatants prepared as previously reported (Sriskandan et al., 1999).

### Human tonsil donors and ethics

Fresh tonsils were obtained by the Imperial College NHS Trust Tissue Bank (ICHTB) from anonymised adult (>16y) donors undergoing routine tonsillectomy and consenting to use of tissue surplus to diagnostic requirement for research. The use of donor tissue was approved by a Research Ethics Committee (07/Q0407/38) and by the Tissue Management Committee of the ICHTB (R12011). Donors (53 females; 12 males) had a median age of 28 years (range 17-59). Indication for tonsillectomy was prolonged or recurrent tonsillitis in all cases bar 4 males with sleep apnoea. Other medical and demographic data were not available.

### Tonsil cell suspensions and histocultures

Tonsil cell suspensions and histocultures were prepared according to previously published methods (Giger et al., 2004). Cells were stimulated at incubation with different concentrations of purified superantigens or bacterial supernatants as above. At the end of experiments, cells were harvested into sterile tubes, centrifuged at 300*g* for 10 minutes, and the supernatant collected and stored at - 20°C. Cells were either used immediately, stored at −80°C in Freezimix (90% foetal calf serum (Invitrogen), 10% DMSO (Sigma) for later flow cytometry analysis. For histocultures, capsule, connective tissue and any damaged tissue was removed from tonsil and remaining tissue dissected into 2-3 mm blocks using a scalpel and placed on 1×1×2cm Gelfoam constructs (Upjohn, Pfizer, UK). Culture supernatants were collected from the Gelfoam constructs at the end of the experiment and stored at −20°C.

### Cellular and immunological assays

Soluble mediators in cell-free tonsil supernatants were measured by ELISA (immunoglobulin EIA, Bethyl Labs USA; cytokine Duosets EIA kits, R&D systems, UK). Tonsil cell proliferation was measured by ^3^H-labelled tritium incorporation and TCRVβ subsets were evaluated by flow cytometry. Human tonsil cells were separated using AutoMACS bead technology (Milltenyi Biotech), either before or after culture. T cells were isolated using CD2 positive selection. B cells were then isolated using the B cell negative selection kit, which positively selected for CD2, CD14, CD16, CD36, CD43 and CD235a, leaving only B cells remaining. Confirmation of the quality of separation was performed using surface staining for CD3 (T cells) and CD20 (B cells), and gave a >99% cell purity on each occasion. Tonsil cell migration in response to CXCL13 was evaluated using the transwell technique as previously described (Debes et al., 2002), using 100nM of CXCL13, a concentration approximately 5 times that secreted by unstimulated tonsil cell cultures.

### Statistics

All statistical analyses, unless stated specifically elsewhere, were performed using Graphpad Prism 5 software. All data were non-parametric; two group comparison Mann-Whitney test was performed, or Wilcoxon signed ranked paired test for paired data. For comparison of multiple groups, a 1 way ANOVA (Kruskal Wallis, or Friedman for paired data) test was performed. Statistical tests were not performed on groups with sample size less than 3. A probability value <5% was classed as significant (p<0.05).

## Results

### Tonsil T cells in suspension cultures proliferate in response to streptococcal superantigens associated with release of pro-inflammatory cytokines

To establish that fresh human tonsil cell suspensions were responsive to streptococcal superantigens, cells were exposed to purified superantigens SpeA and streptococcal mitogenic exotoxin Z (SmeZ), and proliferation assessed by thymidine incorporation (Figure 1-A). Whole human tonsil cell preparations demonstrated dose responsive proliferation similar to that seen with peripheral blood mononuclear cells. Specific superantigen responses were also seen using filter-sterilised bacterial supernatants from isogenic *emm*1 and *emm*89 streptococcal strains differing only in SpeA or SmeZ expression respectively (Figure 1-B). Characteristic expansion of TCRVβ T cell subsets was demonstrated in tonsil cells, most clearly by SmeZ (Supplementary Figure 1) due to the high proportion of TCRVβ8 cells present in unstimulated tonsil preparations.

**Figure 1:**
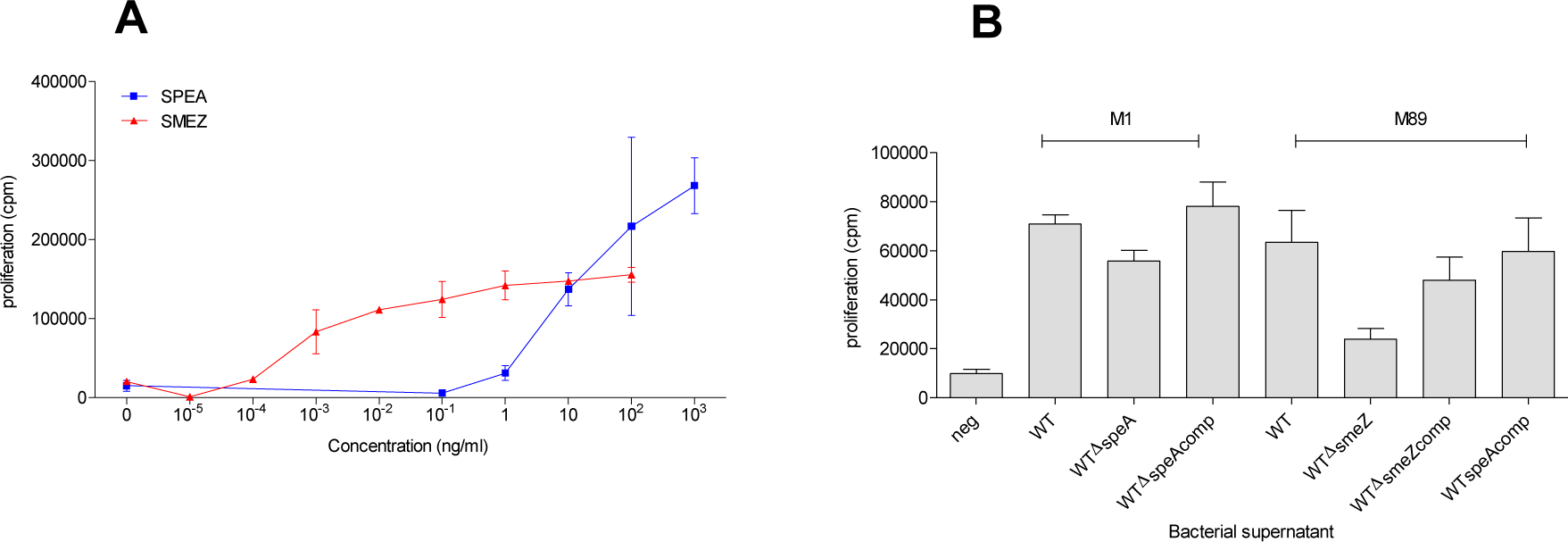
Tonsil cell proliferation in response to purified streptococcal superantigens or streptococcal supernatants. Tonsil cells were stimulated for 7d with different concentrations of purified superantigens (A) or 1% supernatants from isogenic *emm*1 or *emm*89 strains (B). *emm*1 supernatants were from parent (WT); isogenic *speA* negative (WTΔspeA); and complemented (WTΔspeAcomp) strains. *emm*89 supernatants were from parent (WT); isogenic *smeZ* negative (WTΔsmeZ); complemented (WTΔsmeZcomp); and speA complemented (WTspeAcomp) strains. Unstimulated control cells (neg) were incubated with medium alone. Data show the mean and SD from 3 different tonsil donors.

Tonsil cell suspensions stimulated with SpeA demonstrated production of a range of pro-inflammatory cytokines over 7 days, compared to unstimulated controls, where there was minimal cytokine production (Figure 2 and Table 1). Taken together, the findings confirmed that explanted tonsil cell suspensions responded to streptococcal superantigens in a manner similar to previously demonstrated responses in peripheral blood.

**Table 1.** Peak cytokine production in unstimulated and SpeA-stimulated tonsil cells.

| Cytokine | Peak day of production | Median unstimulated (pg/ml) | Median SpeA-stimulated (pg/ml) | Number of tonsil donors | Significance<sup>§</sup> |
| --- | --- | --- | --- | --- | --- |
| IL2 | 2 | 30 | 938 | 5 | p=0.012 |
| IL10 | 4 | 86 | 581 | 5 | p=0.036 |
| IL17 | 4-7 | 567 | 2241 | 7 | p=0.0156 |
| TNFα | 4 | 145 | 1054 | 12 | p=0.0005 |
| TNFβ | 4 | 143 | 3616 | 8 | p=0.0078 |
| INFγ | 5 | 10 | 535 | 7 | p=0.0156 |
| TGFβ | 7 | 129 | 480 | 7 | p=0.0156 |

<sup>§</sup> IL2 and IL10 were assessed by Mann Whitney Test, as numbers were too small for effective pairing; others were assessed using the Wilcoxon matched-pairs signed rank test.

**Figure 2:**
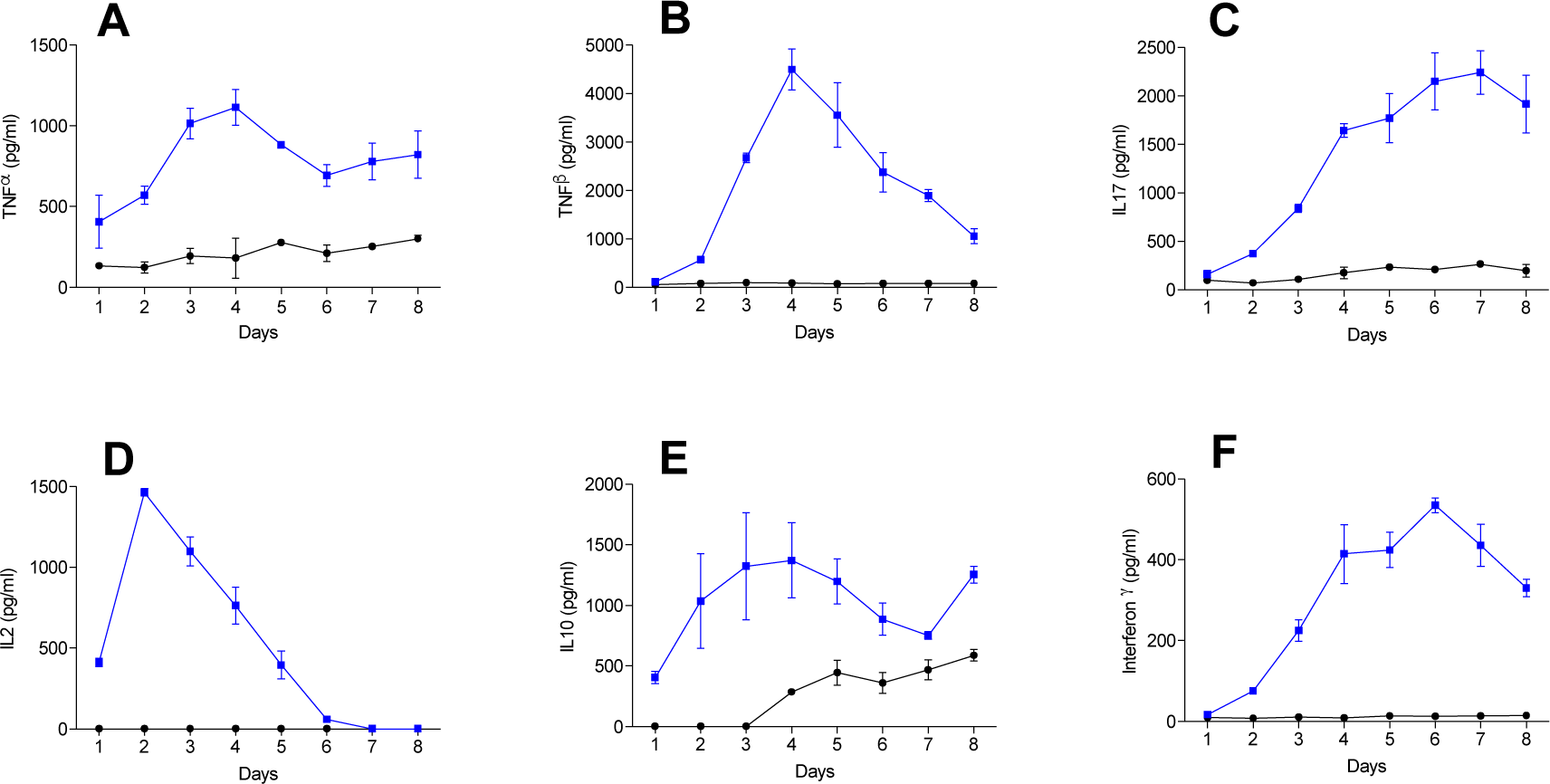
SpeA induces pro-inflammatory cytokine release by stimulated tonsil T cells. Tonsil cells were stimulated for 1 week with recombinant SPEA 100ng/ml (blue) or unstimulated control (black) and supernatants harvested daily from days 1 to 8 of culture. Harvested supernatants were then analysed by ELISA for the presence of cytokines TNFα (A), TNFβ (B), IL17 (C), IL2 (D), IL10 (E) and interferon-γ (F). Mean and standard deviation of experimental triplicates for a single representative donor is shown for each cytokine, representative of between 5-12 different donors measured for each cytokine (Table 1).

### Superantigen stimulation causes a loss of B cells and reduction in total immunoglobulin

Unstimulated tonsil cell suspensions show a mix of predominantly B and T cells, representative of the germinal centres present within tonsils. Stimulation with SpeA caused a profound shift in the cell populations, with a marked loss of B cells compared to unstimulated cells, despite expansion of the T cell population (Figure 3). B cells demonstrated apoptosis, as shown by increased expression of Annexin V and PI, with the change evident after 4-5 days of co-culture with SpeA, coinciding with the earlier observed peak in interferon-γ and TNFβ (lymphotoxin-α) production.

**Figure 3.**
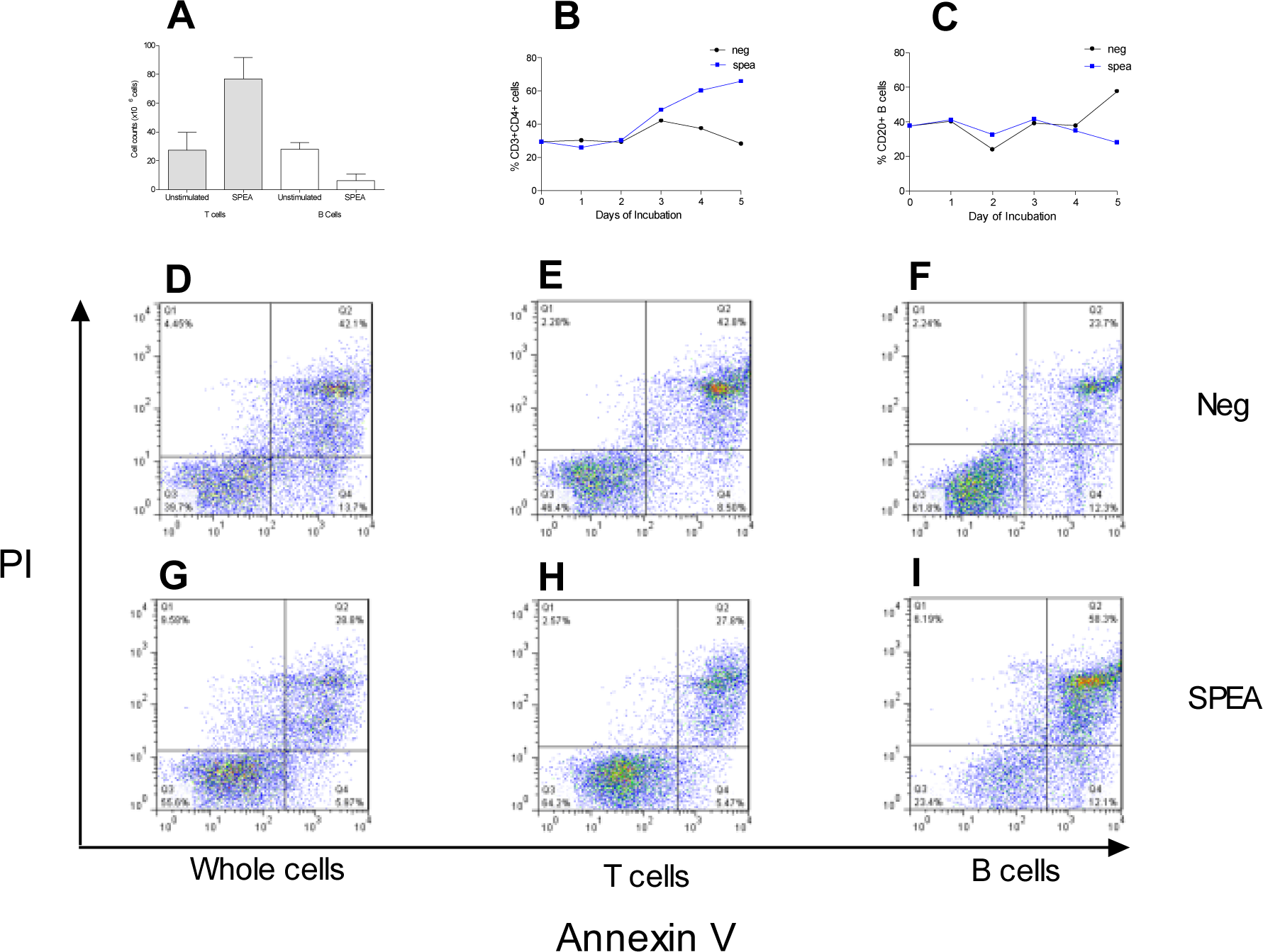
SpeA results in expansion of tonsil T cells but apoptosis of B cells. Tonsil cells were analysed over 5d incubation with SpeA 100ng/ml. Compared to unstimulated controls SpeA resulted in increased total T cell count but a decrease in the total B cell count (A) (N=8 tonsil donors, p=0.0002). The proportions of CD3^+^CD4^+^ tonsil cells (B) increased in the presence of SpeA by day 3 while the reduction in B cells expressing CD20 (C) became apparent by day 5. Annexin V and PI staining of whole cell population showed an overall reduction in apoptosis in SpeA-stimulated cells compared to unstimulated cells (D and G). After cell separation (autoMACS), T cells showed reduced apoptosis in the presence of SpeA (panels E and H), while B cells showed a marked increase in apoptosis (panels F and I), representative plot, N = 2 different tonsil donors.

B cells in unstimulated tonsil cell culture began producing measurably increased quantities of immunoglobulin (IgG, A and M) from day 4 of cell culture (Figure 4). In SpeA-stimulated cultures this response was abrogated, with no increase from baseline. To ensure that this effect was not as a result of tonsil cell processing, tonsils were cultured as histocultures, either unstimulated or treated with SpeA. Histocultures have a shorter survival time in culture, hence responses were measured after 48 hours incubation with SpeA, before onset of tissue necrosis. Tonsil histocultures at 48 hours produced concentrations of IgG that were 2 orders of magnitude greater than tonsil cell suspensions; a reduction in total IgG was again demonstrated in SpeA-stimulated cultures (Figure 4-E).

**Figure 4:**
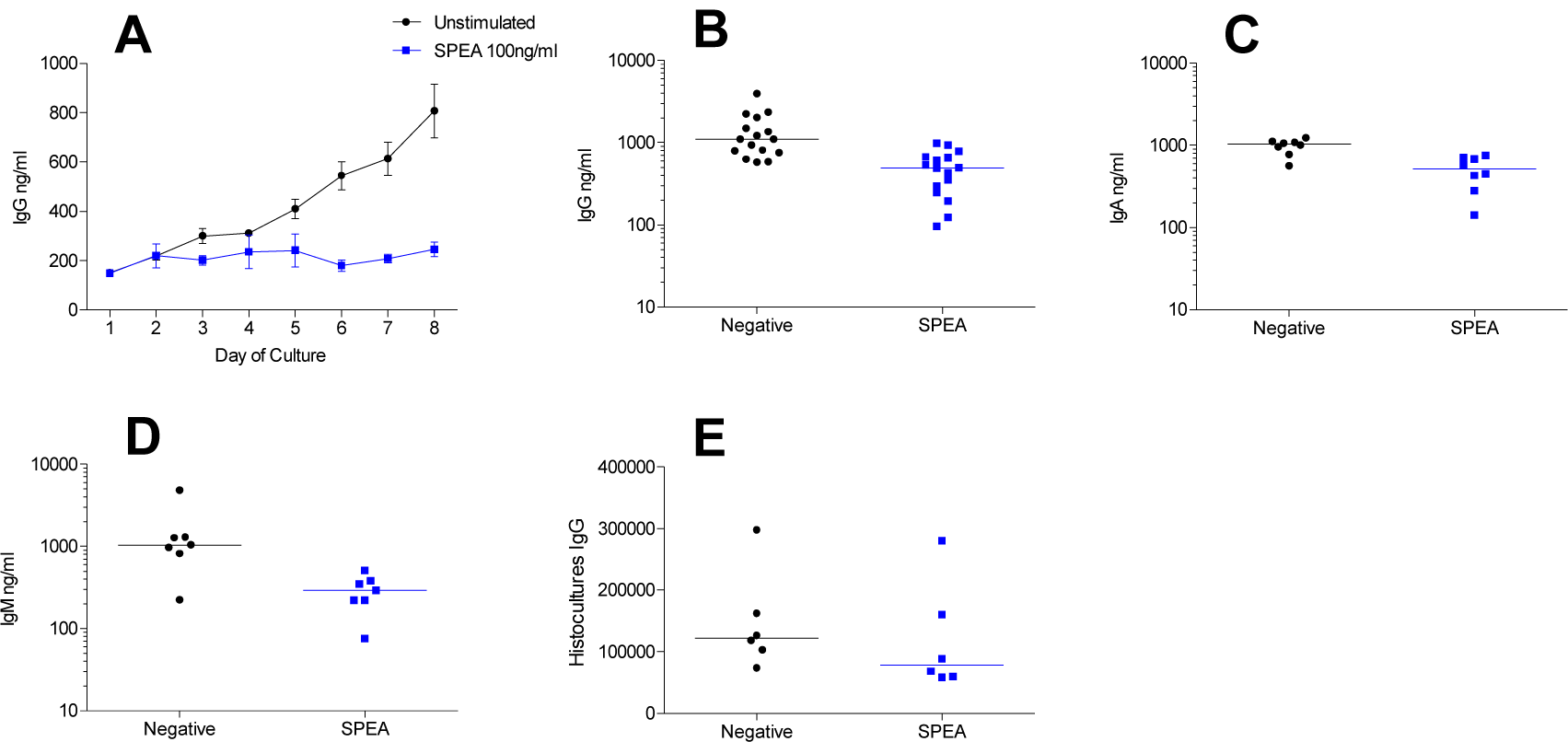
Reduction in tonsil cell immunoglobulin release in response to SpeA. Tonsil cells were stimulated for 1 week with recombinant SpeA 100ng/ml (blue) or unstimulated control (black) and supernatants harvested daily from days 1 to 8 of culture (A). Harvested supernatants were then analysed by ELISA for the presence of total IgG. Means and SD of experimental triplicates from a single donor are shown at each time point, representative of 5 different donors. SpeA significantly reduced tonsil cell production of IgG (B, N=16 donors, p=0.0005); IgA (C, N=8 donors, p=0.0078); and IgM (D, N=7 donors, p=0.0156) in comparison to unstimulated cultures when a wider pool of donors was tested after 7d of co-incubation with SpeA. Horizontal bars represent median values. Tonsil histocultures from 7 donors produced large quantities of IgG (ng/mL) at rest, and this was diminished by co-culture with SpeA (E) p=0.014.

To determine if this effect was due specifically to SpeA, or a result of T cell mitogen-driven proliferation, tonsil cell suspensions were stimulated with a range of known superantigens and mitogens (Figure 5). A reduction in total immunoglobulin was again demonstrated following co-incubation with other streptococcal superantigens, (SpeJ and SmeZ) and staphylococcal superantigens (staphylococcal enterotoxins [SE]B, SEC, and toxic shock syndrome toxin-1 [TSST-1]) in a manner that was dose-dependent (Figure 5A-F). The effect was also demonstrated with the lectin mitogen Concanavalin A (Figure 5G). Stimulation with CD28 antibody produced an increase in immunoglobulin production by tonsil cell suspensions while no change was induced by anti-CD3. However combined anti-CD3 and anti-CD28 resulted in a significant decrease in IgG production by tonsil cells (Figure 5-H). Transferred cytokine-rich supernatants from SpeA-stimulated tonsil cell suspensions into unstimulated cultures did not reproduce the inhibition of total immunoglobulin release (data not shown). The data showed that superantigen-induced T cell proliferation and activation in the tonsil results in B cell apoptosis and loss of function, manifest as immunoglobulin suppression; this effect was observed in tonsil cell suspensions and histocultures and did not appear to be the direct result of any transferrable soluble mediator.

**Figure 5:**
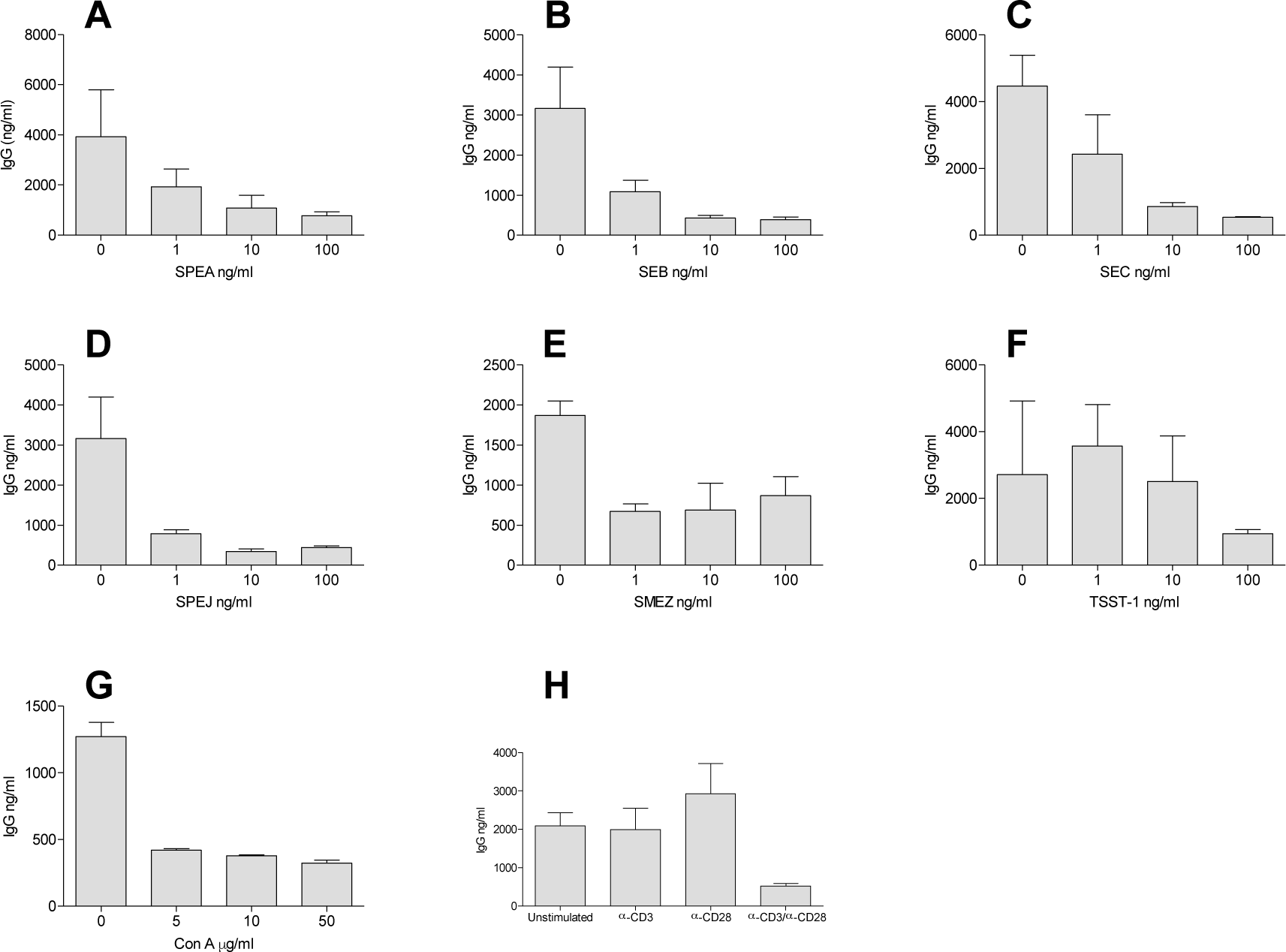
IgG production by tonsil cell suspensions is reduced by superantigens and T cell mitogens. IgG release in the presence of different concentrations of a variety of bacterial superantigens was tested: SpeA (A), SEB (B), SEC (C), SPEJ (D), SmeZ (E) and TSST-1 (F). For comparison, the T cell mitogen concanavalin A was tested (G) and T cell stimulation using anti(α-) CD3 and CD28 antibodies (H). All mitogens inhibited immunoglobulin production in cultures at 1 week. Each graph shows the mean and standard deviation of experimental triplicates from one representative tonsil donor (data representative of experiments using N=3 donors for the different superantigens, N=2 for concanavalin A and α-CD3/28).

### Superantigen phenotype change with SPEA stimulation

T follicular helper (Tfh) cells represent a major proportion of T cells in the tonsil suspensions and are necessary for B cell maturation and function in germinal centres. SpeA induced a significant expansion and increase in expression of key Tfh activation markers OX40 and ICOS (Figure 6A-D). Although the population of CXCR5-expressing T cells expanded in response to SpeA, expression of CXCR5 reduced over time despite increase in other activation markers (Figure 6E and Supplementary Figure 2). To determine if the observed reduction in CXCR5 expression was likely to be functionally relevant, CXCL13 chemotaxis assays were undertaken: SpeA-stimulated tonsil T cells demonstrated reduced chemotactic responses towards CXCL13 compared with controls (Figure 6F).

**Figure 6:**
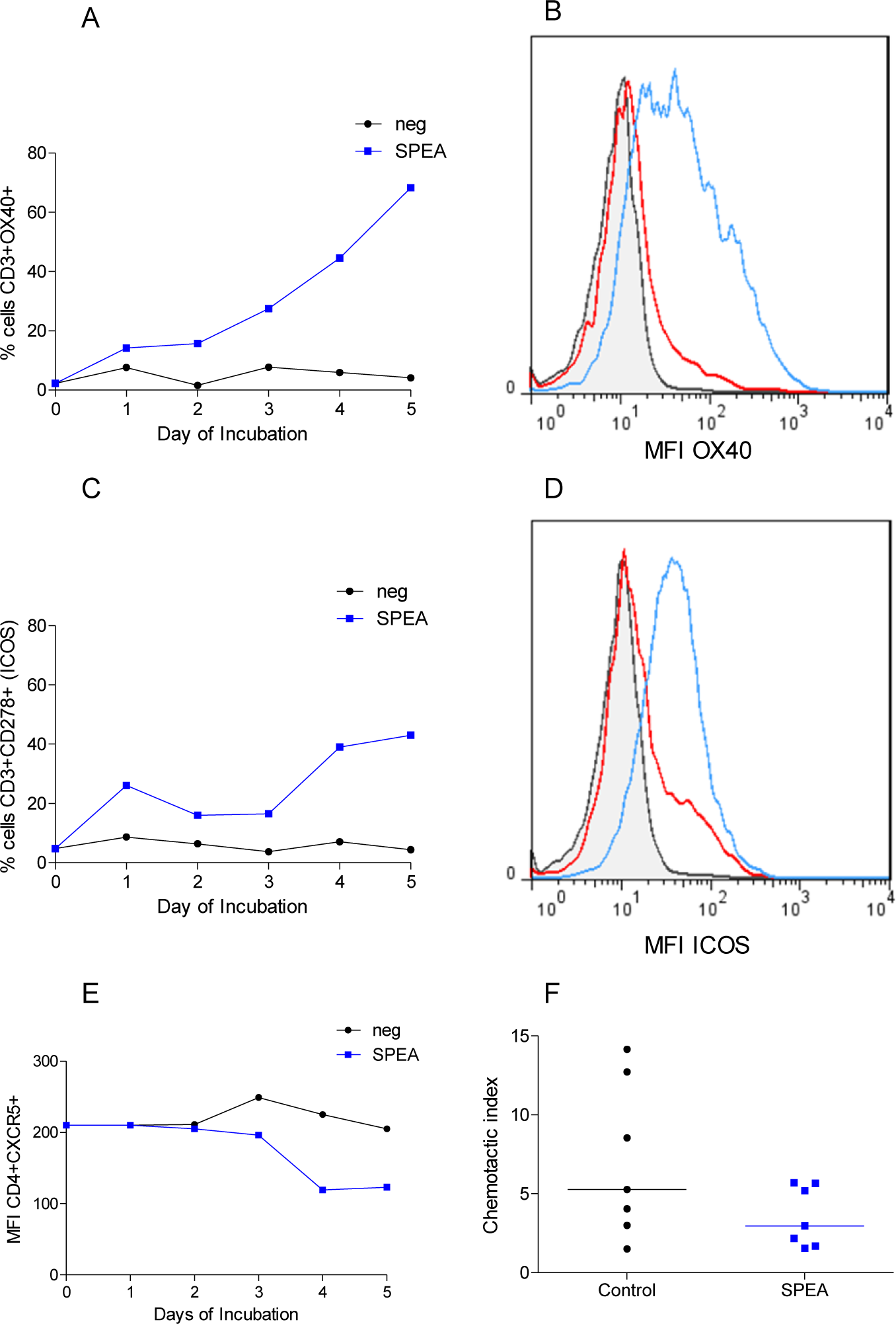
Altered Tfh phenotype in SpeA-stimulated tonsil cells. Tonsil cells stimulated with SpeA (blue) showed a marked early increase in the percentage and mean fluorescence intensity (MFI) of CD3^+^ T cells expressing OX40 (CD134, A and B, p=0.029) and ICOS (CD278, C and D (p=0.029) compared with unstimulated controls (black lines). Representative plots of gated T lymphocytes are shown from 1 donor each; grey shaded, isotype control; red line, unstimulated cells, blue line, SpeA 100ng/ml. In contrast, expression of CXCR5 in CD4^+^ cells reduced in response to SpeA compared with unstimulated controls (E, p=0.015). N=4 tonsil donors for OX40 and ICOS, N=7 for CXCR5. Chemotaxis of tonsil CD4^+^CXCR5^+^ cells toward CXCL13 gradient was reduced by co-incubation with SpeA (F).

## DISCUSSION

This systematic study of tonsil responses to superantigen has demonstrated that *in vitro* primary human tonsil cells demonstrate TCRVβ-specific clonal T cell expansion and pro-inflammatory cytokine release in response to streptococcal superantigens similar to that observed in human peripheral blood mononuclear cells. However, the participating constituent cells in tonsils are markedly different to those in peripheral blood, and cell populations are in close contact. Based in a non-sterile site, the tonsil is repeatedly exposed to bacterial and other antigens; as such, B cells that reside in germinal centres are primed to release significant amounts of immunoglobulin even at rest, supported by a large population of CXCR5-expressing T follicular helper cells that are resident in the tonsil (Crotty, 2011). Focussing on the phage-encoded superantigen SpeA, implicated in the success of the globally successful *emm*/M1 *S. pyogenes* lineage (Nasser et al., 2014; Stevens et al., 1989), we identified a paradoxical reduction in release of multiple classes of immunoglobulin including IgG, IgA and IgM in response to superantigen, despite the marked increase in pro-inflammatory cytokine secretion. Diminution of tonsil immunoglobulin production was reproduced by all superantigens tested, suggesting a common mechanism of immune subversion for both *S. pyogenes* and *Staphylococcus aureus*, and was recapitulated by anti-CD28/anti-CD3. Indeed, a recent report suggested that the presence of superantigen-associated *S. aureus* in tonsil may result in skewing of the tonsil TCRVβ repertoire (Radcliff et al., 2017). Expansion of T cells in tonsil cultures was accompanied by a paradoxical absolute reduction in B cells that was associated with evidence of B cell apoptosis. Tonsil T cell populations expanded by SpeA demonstrated an altered phenotype, expressing high levels of the TNF receptor superfamily members OX40 and ICOS, but paradoxically low levels of CXCR5 compared with unstimulated controls, with reduced chemotactic responses to the chemokine CXCL13. Taken together, the data point to a specific role for streptococcal superantigens in recruiting and diverting tonsil follicular T helper cells from their primary purpose, thereby undermining antibody response during *S. pyogenes* infection, and conceivably other infections that rely on antibody production in the tonsil.

SpeA was first reported to reduce immunoglobulin production almost fifty years ago; a reduction in Ig-secreting B cells was accompanied by a paradoxical increase in spleen weight and 100% expansion in total number of cells in rabbits exposed to SpeA (Hanna & Watson, 1968). Notwithstanding the recognised poor sensitivity of murine cells to superantigens, both SpeA and SpeC were shown to suppress murine spleen B cell IgM responses in a manner that required altered activity of T cells, raising the possibility of either reduced T helper cell function or an increase in T suppressor activity (Cunningham & Watson, 1978). While superantigen-induced T cell-derived secreted products were reported by some to suppress immunoglobulin secreting cells (Bohach et al., 1996; Poindexter & Schlievert, 1986; Proft & Fraser, 2016), others have reported contact-dependent T cell-induced CD95-dependent B cell apoptosis following superantigen exposure in human peripheral blood mononuclear cells. In our study we did not measure CD95L although levels of CD95 were increased. In the tonsil cultures used in the current report, reduction in immunoglobulin required co-incubation of B and T cells and could not be recapitulated by secreted products from T cells; furthermore inhibition of mediators such as IFN-γ or IL-10 did not reverse the observed reduction in immunoglobulin (data not shown), pointing to a requirement for contact between the effector T cells and apoptosing B cells, at least in the experimental setting, and suggesting that altered chemotactic activity may not be the sole mechanism of antibody inhibition.

Within the tonsil and other solid lymphoid organs, distinct populations of T follicular helper cells, characterized by expression of CXCR5, OX40 and PD1, are now recognised to underpin B cell viability through migration to germinal centres in response to CXCL13 (Crotty, 2011). More recently, regulatory T follicular cells, have been described to repress B cell activity and the net balance between the two T cell populations, is believed to underlie B cell activity in lymphoid tissue (Vaccari & Franchini, 2018). Dynamic movements of leukocytes between the circulation and lymphoid tissue can allow ingress of regulatory T follicular cells, although we could not model this in the static culture system used herein. We considered the possibility that exposure to SpeA resulted in a disproportionate expansion of resident regulatory T follicular cells however less than 5% of all tonsil cells in culture expressed FoxP3 (data not shown), therefore we conclude that, in the *ex vivo* setting described in the current work, classically defined regulatory cells play little role. This does not exclude the possibility that superantigen-stimulated regulatory follicular T cells may play a greater role *in vivo*, in lymph nodes draining sites of infection, noting that superantigens are reported to activate regulatory T cells in peripheral blood (Taylor & Llewelyn, 2010). Although not meeting the criteria for regulatory T cells, the phenotype of T cells in SpeA-expanded tonsil cell cultures was significantly and consistently altered such that expression of CXCR5 reduced, measurably impacting on chemotactic function in three of four donors, while other markers of Tfh activation such as ICOS were increased. We speculate that, within the tonsil, reduced chemotactic function may render the Tfh cells unable to migrate to germinal centres and promote B cell antibody production providing one potential explanation for our findings in histocultures. Nonetheless it is clear that the altered phenotype of this T cell population requires further evaluation.

Study of human tonsil cellular responses are necessarily limited by availability of surgical tissue, donor heterogeneity, bacterial contamination of samples, and tissue viability, likely accounting for inter-individual variation (J. Kim et al., 2009). The preponderance of female donors in the current study is unexplained but could represent a bias in selection for surgical referral, or tissue bank consent, factors that could not be controlled for. Nonetheless we did not observe a difference between males and females. Individual clinical history in tonsil donors may influence response to superantigens, particularly as the donors were adults with history of tonsillitis as the leading indication for surgery. Repeated exposure to streptococcal infection might be expected to lead to immunity to toxins such as SpeA with consequent neutralisation of experimentally-added toxin by immunoglobulin secreted by cultured tonsil cells; this was not systematically evaluated in the current work. As many adults have neutralising antibody to SpeA (Proft, T., & Fraser, J. D. 2016) and as tonsil cells produce abundant immunoglobulin, the detection of any effect of SpeA within tonsil cell culture is therefore all the more remarkable, underlining the significance of the effects observed. Indeed, in human peripheral blood it is necessary to use bovine rather than human serum in order to observe reproducible effects due to streptococcal superantigens.

SpeA was selected as a paradigm of streptococcal superantigens as it is strongly associated with the *emm*1 genotype that has caused disease worldwide, and historically is associated with scarlet fever. However, SpeA is less potent in human systems than other streptococcal superantigens such as SpeJ and SmeZ (Unnikrishnan et al., 2002), potentially because SpeA-responsive TCRVβ subsets are smaller, though may alter with stimulus strength (Llewelyn et al., 2006). We used SpeA at a concentration that is physiologically relevant, representing typical concentrations achieved in broth culture (Sriskandan et al., 1999), noting that the concentrations of superantigens within the tonsil have not been characterised.

Since the emergence of invasive group A streptococcal infections in the late 1980’s and description of a superantigen-associated toxic shock-like syndrome (Stevens et al., 1989), there have been a number of reports that report superantigen-related effects in invasive human infection (Sriskandan et al., 1996; Watanabe-Ohnishi et al., 1995). Data showing superantigen-dependent effects in humanised transgenic mice support a role for superantigens in severe disease pathogenesis and inflammation (Proft & Fraser, 2016; Sriskandan et al., 2001), however invasive infection is rare, and the evolutionary benefit of superantigen production is most likely to be manifest in more common non-invasive infections such as tonsillo-pharyngitis. Recent studies in humanised transgenic mice point to a role for SpeA in perpetuating nasopharyngeal infection through a mechanism that requires TCRVβ-specific T cell expansion (Zeppa et al., 2017), while others have highlighted impairment of memory B cell responses to first streptococcal skin infections in mice (Pandey et al., 2016). The extent to which the current findings in human tonsils explain the reported phenomena in mice is unclear, noting that the composition of tonsil is quite different to mucosal and nasal associated lymphoid tissue. Certainly, if superantigens such as SpeA do subvert production of antibody and initiation of adaptive immune response to *S. pyogenes*, this provides a rationale for inclusion of SpeA and other superantigen toxoids in any future *S. pyogenes* vaccine, in order to provide neutralising immunity against SpeA and other superantigens. A similar approach has been proposed for *S. aureus*, where development of natural adaptive immunity may be thwarted by exposure to the B cell superantigen staphylococcal protein A (Kim, et al., 2010).

Leukocytes from the peripheral blood produce limited amounts of immunoglobulin in cell culture; the current work underlines the importance of studying relevant lymphoid tissue when evaluating responses to bacterial pathogens. The human tonsil provides an excellent model system for study of streptococcal disease that can reduce and replace the use of animals in such research, although cannot completely reproduce the dynamic changes in cell populations that may occur in a draining lymph node. Changes in the practice of tonsillectomy are however reducing tissue availability and this will necessitate further refinement of such model systems to reproduce the cell populations found in lymphoid tissue. The current work has identified a clear need to further understand the impact of superantigens such as SpeA on development of adaptive immunity to *S. pyogenes*, elucidation of which is likely to require a nuanced approach combining transgenic mouse models and human cells.

## Acknowledgements

The authors would like to extend their thanks to Mr Shula and his associated surgical team for participation in the ICHT Tissue Bank. FD and SS acknowledge the support of the NIHR Biomedical Research Centre (BRC) awarded to Imperial College NHS Healthcare Trust and the NIHR BRC-supported ICHT Tissue Bank. The authors thank Dr Jennifer Goldblatt for support in chemotaxis assay development and Dr Thomas Proft for reagents.

## Supplementary Information

**Supplementary Table 1:** Flow cytometry antibodies used in this study.

| Analysis group | Purpose | Antibodies |
| --- | --- | --- |
| T cells | Cell characteristics | CD3; CD4; CD8; TCRVβ 2, 8, 11, 14; IOTest TCRVβ repertoire kit; CD25; FoxP3; CXCR5; TCR$\alpha\beta$ |
|  | Activation markers | CD69; CD27; CD278 (ICOS); CD134 (OX40); CD28 |
|  | Apoptosis markers | CD95 |
|  | Immune synapse | CD150 (SLAM); CD154 (CD40L); CD70 |
| B cells | Cell characteristics | CD19; CD20; CD21; CD23; CD27; CD38; HLADR; IgD; IgM |
|  | Activation markers | CD69; CD80; CD86 |
|  | Apoptosis markers | CD95 |
|  | Immune synapse | CD150 (SLAM); OX40L; CD275 (ICOSL); CD40; CD70 |
| Other | Cell type identification | CD14; CD68; CD11b; CD11c; CD56 |
|  | Apoptosis | Annexin V; Propidium iodide |
| | Intracellular cytokines | IL4; IL9; IL17; INF $\gamma$ |

### Supplementary Figures

**Supplementary Figure 1.**
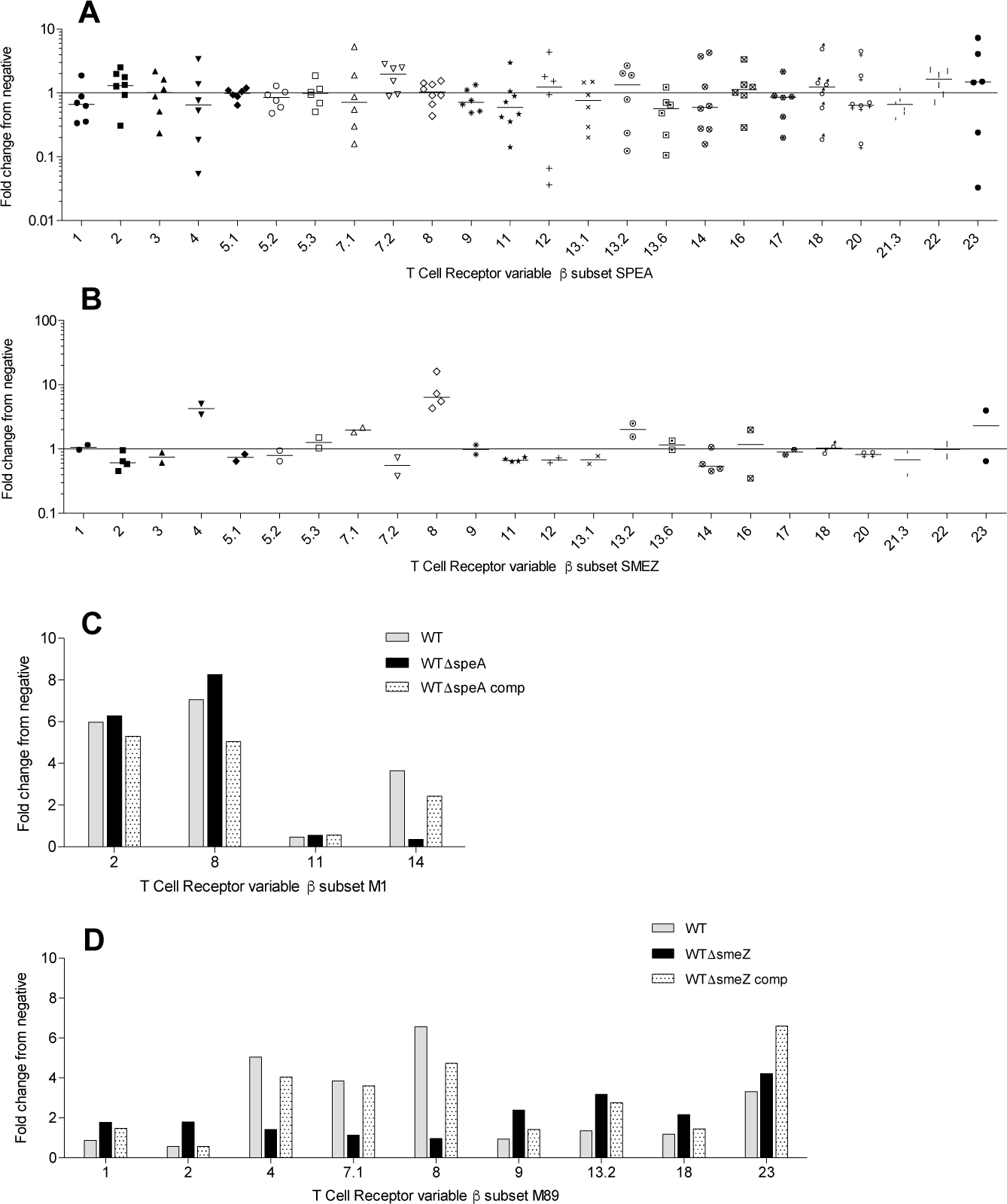
Superantigen-induced tonsil cell TCRVβ profile in different tonsil donors. Fold change from baseline profile is shown following superantigen stimulation for 7d with SpeA (A) and SmeZ (B). Each data point represents a single donor: 6-8 donors (SpeA), 2-4 donors (SmeZ). In two donors, only 4 TCRVβ subsets were checked (Vβ2, 8, 11 and 14). Tonsil cell TCRVβ profile for Vβ2, 8, 11 and 14 following *emm*/M1 streptococcal supernatant stimulation in a single tonsil donor confirmed SpeA-specific TCRVβ changes (C) when comparing supernatants from parent (WT); isogenic *speA* negative (WTΔspeA); and complemented (WTΔspeA comp) strains with unstimulated cells (neg) from the same tonsil donor. Tonsil cell TCRVβ profile following *emm*/M89 supernatant stimulation in a single tonsil donor confirmed SmeZ-specific changes (D) when comparing supernatants from parent (WT); isogenic *smeZ* negative (WTΔsmeZ); and complemented (WTΔsmeZ comp) strains with unstimulated cultures from the same tonsil donor. Results for the TCRVβ subsets 1, 2, 4, 7.1, 8, 9, 13.2, 18 and 23 only are shown, as there was no alteration from baseline with the other TCRVβ subsets tested.

**Supplementary Figure 2.**
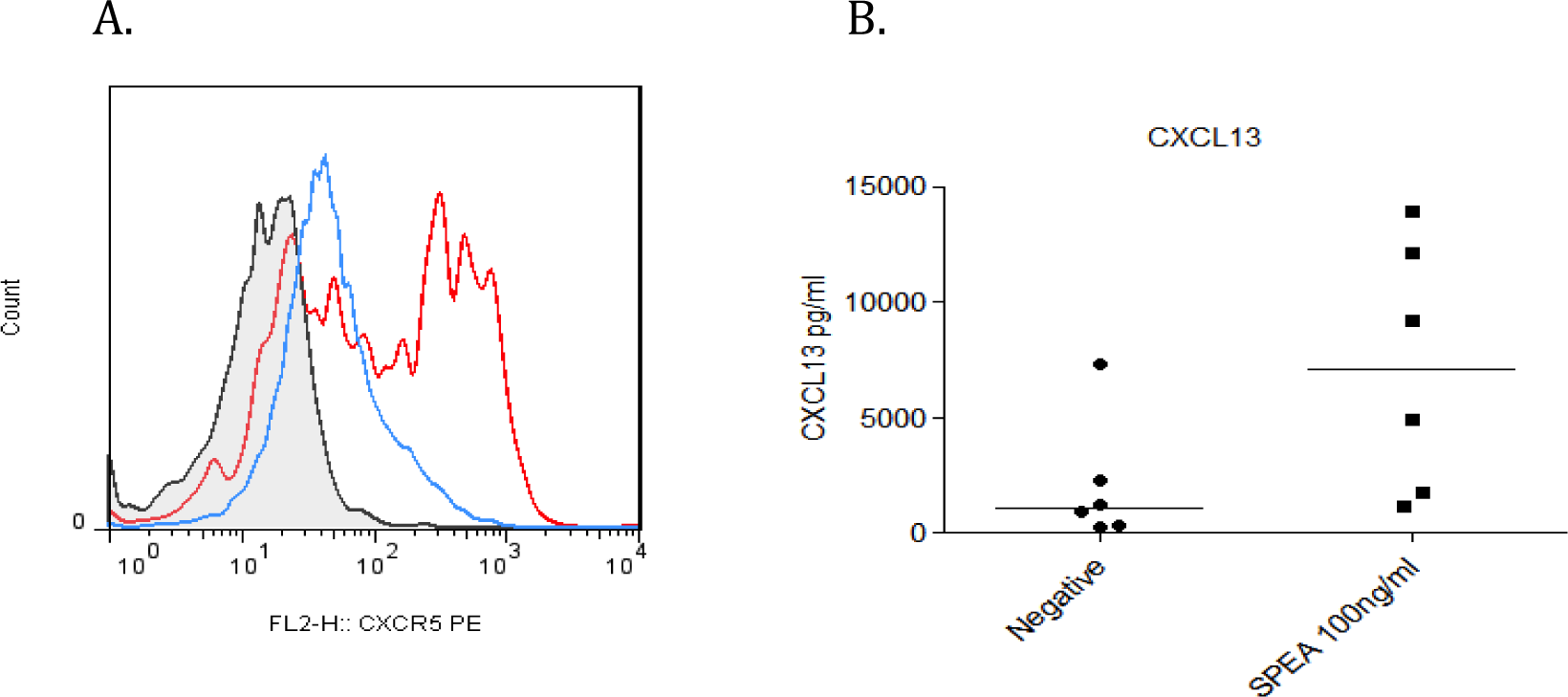
Reduction in CXCR5 expression within tonsil T cell population in response to SpeA stimulation despite increased CXCL13 release. Mean fluorescence intensity shown for CXCR5+ gated tonsil T cells (A) Blue line, SpeA; Red line, unstimulated controls; Black line, isotype control antibody. Plot from single donor representative of 7 tested donors. (B) CXCL13 levels in SpeA-stimulated and control tonsil cell suspensions

